# PeerPub: A device for concurrent operant oral self-administration by multiple rats

**DOI:** 10.1101/2022.03.25.485830

**Authors:** Paige M. Lemen, Jie Ni, Jun Huang, Hao Chen

## Abstract

The social environment has long been recognized to play an important role in substance abuse disorders (SUD). Operant conditioning is the most widely used rodent model of SUD. However, most operant chambers do not accommodate more than one rat at a time. Here, we introduce PeerPub – a novel social operant chamber. PeerPub uses a touch sensor to record the number of licks on drinking spouts. It then delivers a drop of solution with a fixed volume as the reward to the tip of the spout when the number of licks meets the requirement of a reinforcement schedule. A radio-frequency identification (RFID) chip implanted on top of each rat’s skull tracks the identity of the rat. The system is controlled by a Raspberry Pi computer. We tested PeerPub using male Wistar Kyoto rats in daily one-hour sessions where supersac, a solution containing glucose and saccharin, was delivered under a fixed ratio 5 schedule. We found that male rats consumed more supersac in group housing rather than in isolated conditions. These data demonstrated the utility of PeerPub in modeling the interaction between motivated behavior and social context. We anticipate devices like PeerPub will help demonstrate the role of the social environment in SUD phenotypes. The design of PeerPub is available at http://github.com/nijie321/PeerPub

## Introduction

There have been many groundbreaking studies on mechanisms of substance abuse disorder (SUD), but translating these research findings into prevention and treatment strategies has been very challenging (Heilig et al., 2016; Venniro et al., 2020). One potential explanation is that many environmental factors that influence SUD, such as adverse childhood experiences, chronic exposure to stress, and the role of social environment, are not incorporated into studies using model organisms. Operant intravenous drug self-administration is one of the most widely used paradigms to model SUD. It allows an animal to acquire voluntary drug consumption by learning the association between an action (e.g., lever pressing, nose poking, or spout licking) with the delivery of a reward. Most operant conditioning experiments were conducted using operant chambers that completely isolate each individual, which precluded the role of social environment on drug-taking behavior to be studied. A much smaller number of studies, however, have investigated the role of the social environment in SUD using rodent models. For example, it has been reported that rats tested with a drug-experienced partner were faster to acquire cocaine self-administration (Gipson et al., 2011; Smith et al., 2014), and social learning facilitated nicotine self-administration (Chen et al., 2011). Although physical isolation is required to preserve the intravenous catheters used for drug delivery, these studies used specially constructed operant chambers that isolated individual rats but allowed limited social contact via either a mesh (Peartree et al., 2017) or several small windows (Chen et al., 2011).

Voluntary oral consumption of drugs provides an opportunity to model SUD using rodents in more realistic social settings. For example, the Rat Park set of experiments found that rats living in a social environment were less likely to self-administer morphine orally, as opposed to rats living in an isolated environment (Alexander et al., 1978). These experiments were conducted using a device that recorded videos of rats during drug consumption, and each rat was identified by the hair dye mark on their back (Coambs et al., 1980). These pioneering studies, however, did not use operant conditioning procedures.

Here, we describe PeerPub – a device that allows multiple rats to learn operant oral drug self-administration. PeerPub used touch sensors to record licking events. These events were assigned to individual rats based on the unique radio-frequency signal detected from a microchip embedded on top of each rat’s skull at the time of lick. A drop of solution was delivered to the tip of the spout when the number of licks meets the criteria of a reinforcement schedule.

We tested PeerPub using Sprague Dawley rats by allowing them access to Supersac (a solution containing glucose and saccharin) under the control of a fixed ratio (FR) 5 schedule. We found a significant interaction between social environment and sex on supersac oral self-administration: males self-administered significantly more supersac in social than in isolated environment, while females obtained similar amounts of supersac in both environments.

## Methods

### Overview of the system

PeerPub is a novel operant chamber that records the operant licking behavior of multiple (we tested two) rats simultaneously (Figure 1). Each PeerPub device has two stainless steel rodent lickets (with a stainless steel ball at the end). One of the spouts is designated to be active and the other one is inactive. These spouts are installed in two identical 3D printed spout holders. Each holder connects the spout to a unique channel on a capacitive touch sensor, which records the timing of each lick with millisecond time resolution. The faceplate of the spout holder contains an antenna for the radio-frequency identification (RFID) system. Each rat has an RFID chip embedded on top of the skull. Each lick is assigned to the RFID detected at the time of licking. Upon the completion of a reinforcement schedule on the active spout, a drop of solution (i.e., the reward) is pushed to the tip of the spout via a syringe driven by a step motor (i.e., a pump). Licking on the inactive spout has no consequences but all the licks are recorded. The system is controlled by a Raspberry Pi (RPi) computer. A set of RFID fobs with their IDs stored in the software is used to set the session length and reinforcement schedule (e.g., fixed or variable ratios) and start each test session. Data are transferred wirelessly to a remote server upon the completion of each session. We did not include an operant chamber in our design. Both the spout holders and the case of the RPi are 77 mm in width. They can be installed in standard rat operant chambers available from several commercial sources.

**Figure 1:**
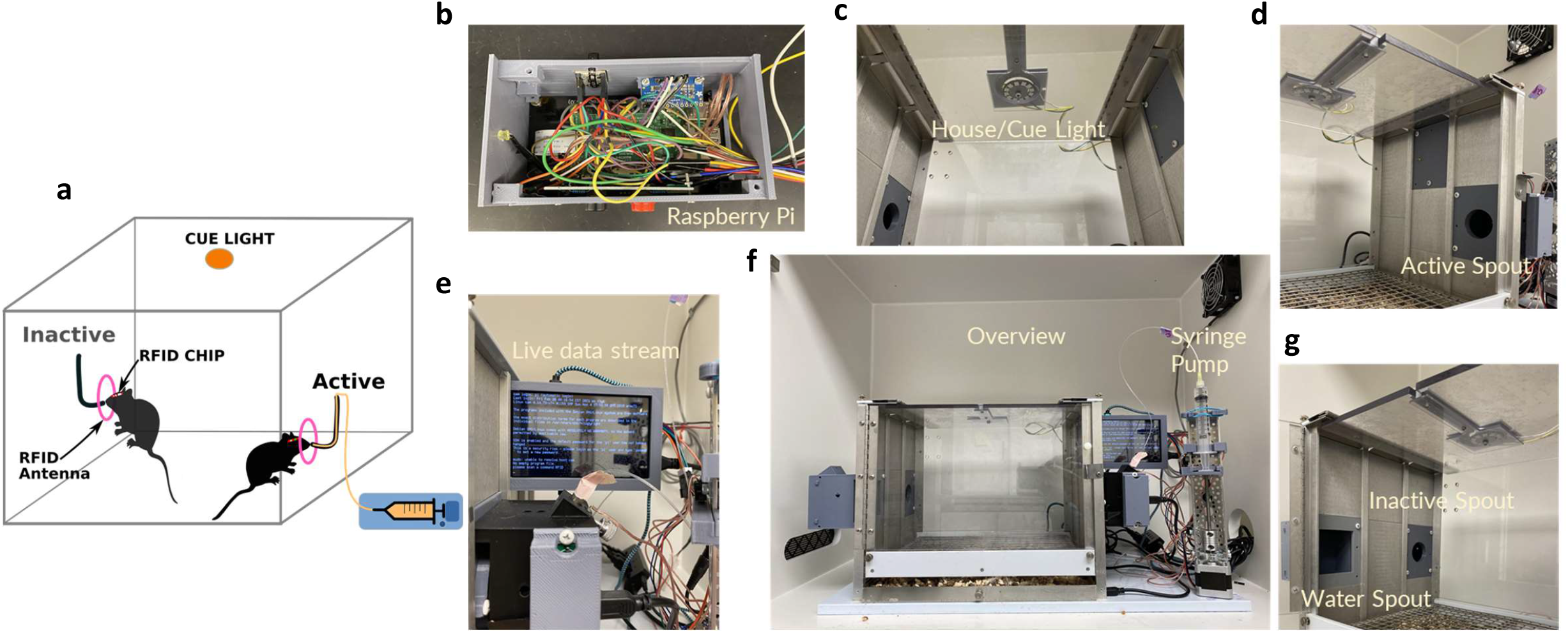
(a) Drug self-administration chamber set-up for recording drug intake and licking microstructure of 2 rats simultaneously. (b) The raspberry Pi computer that collects data in real time, place on the side of the chamber. (c) The cue light that flashes each time a reward is given by the pump. (d) The “active” spout and holder that is attached to a syringe and pump containing a substance for oral self-administration. (e) The screen of a Raspberry Pi computer attached to the chamber. This shows data being collected in real time. (F) Full set-up of all parts of oral drug self-administration chamber and attached parts. (g) The “inactive” spout is the control spout used to compare licks between spouts with and without drug. Water spout is an optional addition for sessions that run for a long time and animals need access to water. The spout holder is removable and can be replaced with more chamber wall.

### Hardware

The RPi is a credit card-sized single-board computer originally made for electronic hobbyists. We used the RPi 3 (Model B) in this project. Each RPi has 40 general-purpose input/output (GPIO) pins. These pins are the interface between the RPi and other electronic modules, such as touch sensors (input), LED lights (output), step motors (output).

We used the EM4100 RFID system because of the availability of inexpensive readers with sensitivity sufficient for our needs. The injectable RFID chip was 2 × 8 mm, each emitting a unique hexadecimal code when placed in the vicinity of the antenna. The antenna guarding the active spout was configured to output the encoded unique ID as an 8 digit hexadecimal code, while the inactive antenna produced a 10 digit hexadecimal code. This configuration was accomplished using Windows software provided by the manufacturer. This setup allowed the program to distinguish licking behavior emitted from spouts The diameter of the antenna is 45 mm. The antenna is placed vertically behind the spout holder’s faceplate, forcing the rat to place its head inside the antenna loop when licking the spout. We further adjusted the opening of the faceplate to ensure that the rat’s skull is almost touching the upper edge of the opening, where the RFID on the skull will generate the strongest signal. In addition, a set of RFID chips embedded in plastic fobs were used to start each session with specific configurations. These fobs allowed lab personnel to start the session without using a keyboard.

A NeoPixel RGB LED Ring is a small electronic component that consists of addressable RGB LEDs arranged in a closely spaced circle and each LED can be controlled by outputting a tuple of three 8 bit values ranging from 0 to 255 to control color and brightness. We use a NeoPixel Ring with 12 LEDs as house and cue lights. The House light is turned on for a time period between 9:00 PM to 9:00 AM and turns off the rest of the time (our rats are housed in reverse light cycled rooms). The LED Ring was programmed to flash for 0.5 seconds when a reward was delivered. This acted as a cue for drug reward.

A stepper motor was attached to a rectangular metal channel, along with several 3D printed mount pieces for the syringe tube, for positioning purposes. A stepper motor driver connects the RPi and the stepper motor to allow the configuration of precise steps and rotation. Our pump design was based on the open-source design by Varnon (“OpenSourceSyringePump —,” n.d.). We used two push buttons (forward and backward) connected to the RPi via GPIO to allow manual syringe pump adjustment. After each session the plunger will be at a higher position; it is necessary to reposition it so the syringe can be reloaded before a new session. In addition, two limit buttons were mounted to the side of the channel for the purpose of plunger repositioning to detect that the syringe has reached the end position. We programmed the software to send an alert message via the wireless network to a communication application (Slack) when this happened so that the syringe can be refilled as needed. The forward button will advance the syringe while it is being held in place. In contrast, the back button will automatically lower the plunger until the edge of the plunger hits the limit switch.

We used the Adafruit Capacitive Touch Sensor Breakout (MPR121) to record licking activities. The active and inactive spouts were connected via separate electric wires to the sensor. We only use 2 of the 12 available channels. A 5-inch display was connected to the RPi for visual interpretation and debugging. To fit the case on the side of the chamber, we designed a 3D printed case to fit the RPi and other electronic components connected to the GPIO pins. The case was designed to fit in the slots of standard commercial rat operant chambers but the design can be easily adjusted. We connected the RFID readers to a powered external USB hub to ensure sufficient sensitivity. A 3 Amp power adapter was used to power the RPi.

### Software

We programmed the RPi with logic specific to our experiment. We used Python 3 as the programming language because it supports different third-party libraries for GPIO programming. We used the gpiozero and RPi.GPIO library to control external input/output modules. The main program asks the user to enter a specific type of session, which can be provided by scanning the specific RFID fobs without using a keyboard. The RFIDs implanted in the rats are then scanned. The main program then spawns a separate thread to handle licking behavior logics while the main thread is kept alive to record rats’ RFIDs when they poke their head into the spout holder. The second thread needs the rat’s RFID to keep a count of how many times active and inactive spouts are licked and give out rewards accordingly. The program was written in an object-oriented fashion to keep the code clean. All code, including 3D design, are available at http://github.com/nijie321/PeerPub

We used the Raspbian operating system without a graphical user interface to reduce computational overhead. All operations are done on the console and colored prompt texts are used to provide behavior data in real-time.

### Experimental design

We examined the effect of social isolation housing and supersac self-administration using young adult Sprague Dawley rats (Postnatal day approximately 55 at the onset of the experiment) placed in either a cage alone or group-housed with one other cagemate (Males=10-12/group, Females=8-10/group). These rats went through a 10-day familiarization phase to make sure they were well acquainted with their cagemate or being isolated before beginning our self-administration protocol of 4 hr water deprivation then 1 hr oral supersac self-administration in our chambers attached to PeerPub for 8 sessions (1 water deprivation/session per day). Then, both housing and chamber social status were switched; isolated rats were paired together and the group-housed rats were separated for another 8 sessions (16 sessions total). For this specific experiment, we chose a fixed ratio of 5 (FR5) as the reinforcement schedule. Supersac is a glucose-saccharin solution: 3% glucose + 0.0125% saccharin in water, and its use for measuring self-administration has been an established model for measuring consumption in previous research (June and Gilpin, 2010). Total number of rewards, licks on the active and inactive spurs were recorded for each rat. Body weights were recorded before each session.

## Results

For supersac oral intake, both male and female Sprague Dawley rats showed an intake difference in isolated vs. group-housed and chamber environments, though the male intake difference between groups is more apparent (Figure 2). Male rats that were group housed, and underwent self-administration with their cagemate, took more supersac than the isolated group that was in the chamber undergoing self-administration alone. In females, the difference was minimal but the trend is opposite of males; isolated rats drink more supersac than group housed rats. A one-way repeated measures ANOVA examined the effects of the treatment group (isolated vs. group housed) on intake. Results show that the type of treatment group led to differences in intake for both males (treatment F(1,19)=11.5, p=0.003) and females (treatment F(1,15)=0.14, p=0.71), but only males were statistically significant. Regardless, these results indicate that housing and chamber social environments play a role in intake. After session 8, rats switched groups: previously isolated rats became group housed and previously group housed rats became isolated. Chamber social environment continued to match housing groups. After the switch, supersac intake no longer showed an intake difference between groups in either sex (p>0.05 for all).

**Figure 2:**
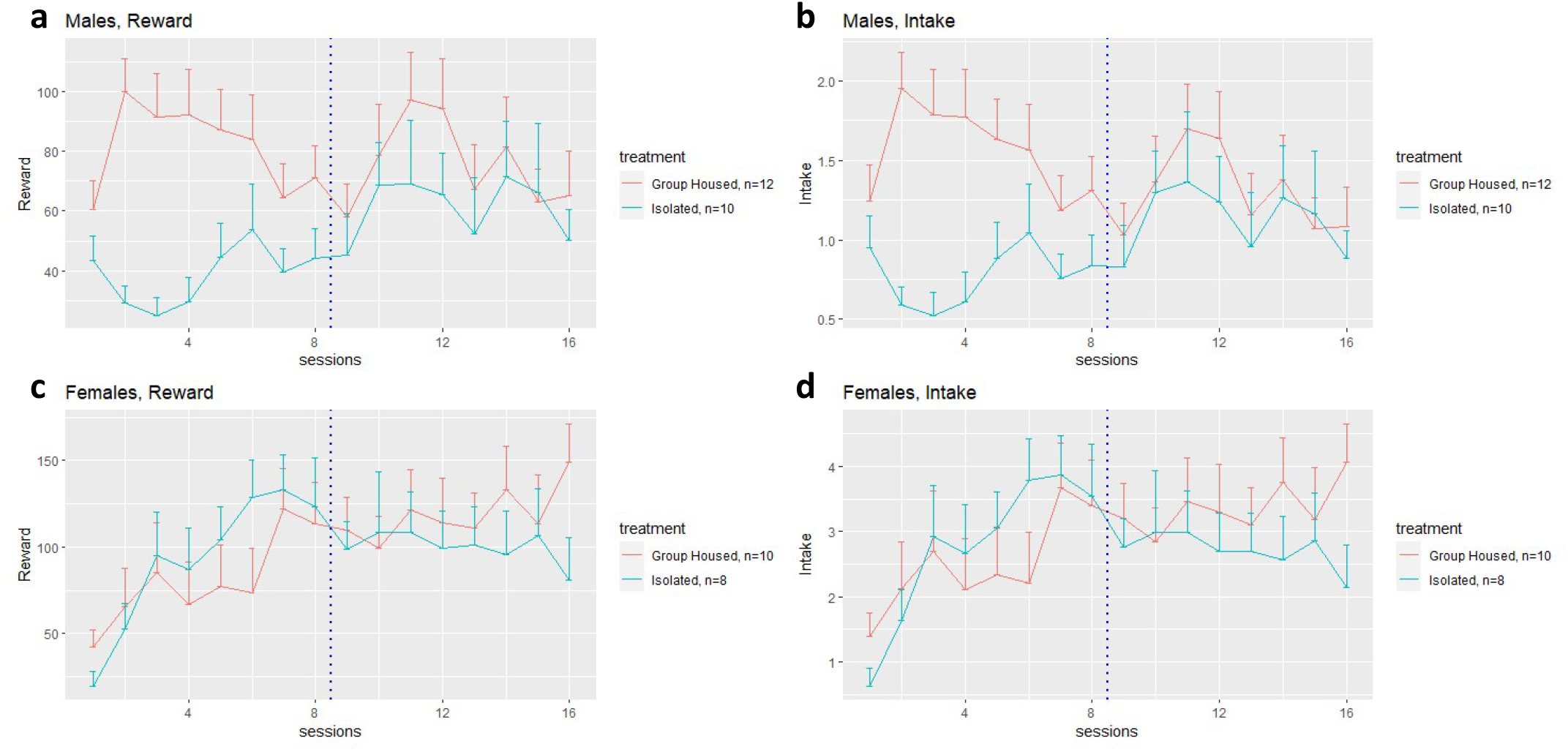
Reward and intake between groups for each sex. Treatment group indicates the group that rats started in. The dotted blue line indicates the switch of group; rats previously group housed are now isolated and rats previously isolated are now group housed. Housing and chamber social environment remained matched. (a) Male number of rewards given per session, group housed vs. isolated. (b) Male intake amount per session, group housed vs isolated. (c) Female number of rewards given per session, group housed vs. isolated. (d) Female intake amount per session, group housed vs isolated.

## Discussion

We designed a novel operant chamber, PeerPub, that monitors the licking behavior of individual rats based on the RFID implanted on the top of their skulls. Licking events triggered the delivery of an appetitive solution upon the completion of a reinforcement schedule. Using the device, we found that male Sprague Dawley rats tested with a peer drank significantly larger amounts of supersac than those tested alone. Females did not show a statistically significant effect of social setting.

The social environment affects food intake. Humans tend to conform eating habits with others as rewarding adaptive behavior (Higgs and Thomas, 2016); we are more likely to follow what is perceived as normal based on social comparison. In Wistar rats, both males and females drink and eat more when previously placed in crowded housing for 6 hrs versus when previously isolated (Brown and Grunberg, 1996). Also, during an 18 hr time frame rats consumed more water and food in crowded housing vs isolated (Brown and Grunberg, 1996). However, there is also evidence showing the opposite. Sprague Dawley rats isolated have been shown to have higher food intake than their socially housed peers (Schipper et al., 2018).

The social environment also affects drug intake. In humans, social influences, such as family, schools, peer groups, and religion, play a role in the likelihood of initial and continued use of substances with the potential of abuse, as well as the likelihood individuals will develop a SUD (Dew et al., 2007). Other social influences, such as socioeconomic status and access to safe community resources, treatment, and support groups, also play a role in drug use and the potential for abuse (Mennis et al., 2016).

Mirroring human behavior for drug addiction research has been historically difficult. Past animal models used for the assessment of substance abuse risk often involve physical stressors, such as restraints, forced swim, etc. (Finnell et al., 2017; Heilig et al., 2016), but this does not model the social determinants that contribute to human addiction risk (Miczek et al., 2008; Yap and Miczek, 2008). Also, animal models for drug abuse often only record one rat for drug intake (Peitz et al., 2013), which has been criticized because it does not represent the social complexities of human life (Heilig et al., 2016; Smith, 2012). Studying an individual animal in isolation may not capture the same behavior as studying a group of animals during drug use; one of the most reliable predictors of whether or not adolescent or young adults will take drugs is whether their friends take drugs (Bahr et al., 2005; Simons-Morton and Chen, 2006; Smith, 2012). PeerPub introduces the possibility of recording multiple rats during drug self-administration. Additionally, we designed PeerPub to record oral drug self-administration because most people with OUD begin using opioids orally before progressing to an injecting or snorting method of drug intake (Katz et al., 2011; National Academies of Sciences, Engineering, and Medicine et al., 2017; Surratt et al., 2011).

Measuring individual rat oral intake in a social setting is possible when using a RPi computer to customize operant chambers for specific experiment requirements. This ability improves on our currently available tools to incorporate the so(Katz et al., 2011; National Academies of Sciences, Engineering, and Medicine et al., 2017; Surratt et al., 2011)cial aspect of food and drug consumption. All current models provide important analysis on substance use, but are still very limited. Previous techniques to study social environments in substance intake involve, for example, separating rats with a wall so that social interaction is only available visually (Gipson et al., 2011), only one rat has access to a spout or lever(June and Gilpin, 2010), or rats are given a choice between drug reward or social reward rather than evaluating intake with both options simultaneously (Venniro and Shaham, 2020).

When animal models undergo oral self-administration, the speed at which they lick the spout is an indication of the subjective value of the reward, i.e, how much they “like” it.. Faster is seen when more appetitive rewards were provided and vice versa. The speed at which rats lick therefore can be used to indicate the subjective experience the substance provides. PeerPub records each lick with a precise timestamp. These data can then be used to analyze the microstructure of these licking bouts.

This method also allows for the study of oral self-administration intake, and is, to the best of our knowledge, the only oral operant chamber for self-administration that measures two rats simultaneously in one chamber. Previous techniques have used levers to signal a reward given intravenously through a catheter implanted surgically to the rat’s jugular vein(Peitz et al., 2013). This kind of surgery is more stressful to the rat and time-consuming compared to implanting an RFID underneath the skin in PeerPub. Catheter surgery also increases the risk of infection and can sometimes be difficult to maintain position. Our method provides a less invasive and risky preparation that can be seen as an improvement for reducing the risk of extra variables, such as stress effects, to consider, as well as improving animal welfare.

While using customizable devices like RPi allow you to have full control in designing experiments, there are still some drawbacks that need to be considered beforehand. Basic computer engineering skills (programming and soldering) are required to modify or add on to the current design, limiting the potential use of this technology to only those that meet the skill requirements.

Based on these results, it is clear that the current social environment in housing and self-administration chambers plays a role in intake. Additionally, the results shown after switching rats’ social environment indicate that changes in the social environment likely affect the oral intake of substances and, therefore, is important to study its role in drug use when using animal models to better represent human social factors that play a role in drug consumption.

PeerPub provides a new way to evaluate the motivation of oral drug consumption in a social setting by recording multiple rats simultaneously in the same space. It provides a new avenue to studying the role of the social environment in the use and misuse of substances.

